# ADAM10 mediates macroglial cell fate decisions in the developing brain

**DOI:** 10.1101/2023.02.11.527059

**Authors:** Yihui Wang, Yue Liang, Daosheng Ai, Jun-Liszt Li, Ziyan Feng, Zhibao Guo, Xing-jun Chen, Tingting Zhang, Xiaoxiao Zou, Jun-Li Gao, Xiaofei Gao, Xiao-Ling Hu, Long-Jun Wu, Wenzhi Sun, Suiqiang Zhu, Shumin Duan, Wei Wang, Woo-ping Ge

## Abstract

ADAMs (<u>a d</u>isintegrin <u>a</u>nd <u>m</u>etalloproteinase) are transmembrane proteins with cell adhesion and protease activities that contain disintegrin and metalloproteinase domains. ADAM10, a member of the ADAM family, is widely expressed in the brain. There are >40 substrates reported for ADAM10, including Notch, Delta-like ligand-1 (Dll1), and N-cadherin. To date, however, its function in the brain has been largely unknown. We used genetic manipulation to delete *Adam10* specifically from glial progenitors in developing brains and observed that conditional knockout mice showed locomotor abnormalities. They all died within 4 months with apparent defects in the cerebellum. By comprehensively analyzing data from bulk RNA sequencing, single-cell RNA sequencing, and staining of the cerebellum, we found that ADAM10 promoted astrocyte generation under physiological conditions. Upon the removal of *Adam10* in glial progenitors, the production of oligodendrocytes vastly increased, whereas the generation of astrocytes was substantially inhibited. Our results showed that ADAM10 plays a critical role in macroglial cell fate decisions during brain development.

## Introduction

Glial cells play a crucial role in the development, metabolism, homeostasis, and function of the brain. Astrocytes and oligodendrocytes, two major macroglial populations derived initially from neuroepithelial progenitor cells, are widely distributed throughout the central nervous system (CNS)(Rowitch and Kriegstein, 2010). With numerous processes enwrapping synapses, astrocytes shape neural circuits during their development and subsequently help to maintain their functions(Chung et al., 2015; Chung and Barres, 2012; Freeman, 2015). Oligodendrocytes form myelin sheaths that wrap around neuronal axons(Freeman and Rowitch, 2013). They promote the rapid transmission of action potentials and provide metabolic support for the wrapped axons(Rowitch and Kriegstein, 2010). The dysregulation of oligodendrocyte development leads to neuronal death in psychiatric disorders(Fields, 2008) and multiple neurodegenerative diseases, including multiple sclerosis (MS) and amyotrophic lateral sclerosis (ALS)(Saab et al., 2013).

The generation of neurons and glia in the cerebral cortex follows an intrinsic developmental sequence, in which neurons are produced first, followed by astrocytes and oligodendrocytes(Qian et al., 2000). Under strict regulation, the sequential appearance of neurons and glia in the brain is controlled by competition among many growth-factor signaling pathways and downstream transcription factors(Sauvageot and Stiles, 2002). For example, platelet-derived growth factor (PDGF) drives neuronal differentiation(Williams et al., 1997), whereas cytokines (e.g., ciliary neurotrophic factor [CNTF], interleukin 6 [IL-6], and leukemia inhibitory factor [LIF]) and Notch family members promote astrocytic fate determination(Barnabé-Heider et al., 2005; Bonni et al., 1997; Gaiano et al., 2000; Rajan and McKay, 1998). Activation of the Notch signaling pathway promotes gliogenesis(Irvin et al., 2001; Tanigaki et al., 2001). The bHLH transcription factor neurogenin (Ngn) actively participates in neurogenesis but inhibits the differentiation of neural stem cells (NSCs) into astrocytes(Nieto et al., 2001; Sun et al., 2001). However, the bone morphogenetic proteins (BMPs) promote either neuronal or astrocytic fates depending on the age of the cell(Nakashima et al., 2001). The Janus kinase/signal transducers and activators of transcription (JAK/STAT) signaling pathway responds to CNTF-driven astrocyte formation(Bonni *et al*., 1997; Nakashima et al., 1999). Oligodendrocyte precursor cells are derived from radial glia and then differentiate into mature oligodendrocytes(Freeman and Rowitch, 2013). Sonic hedgehog (Shh) drives oligodendrocyte formation by altering the expression of transcription factors(Alberta et al., 2001; Tekki-Kessaris et al., 2001). Both oligodendrocyte transcription factor 1/2 (Olig1/2) and SRY-box transcription factor 10 (Sox10) are involved in oligodendrocyte development and work together to determine the identity of oligodendrocytes(Lu et al., 2001; Sock and Wegner, 2021; Zhou et al., 2000).

ADAMs are a family of transmembrane proteins with disintegrin and metalloproteinase domains. There are 29 members in the ADAM family(Edwards et al., 2008; Primakoff and Myles, 2000). These proteins have both cell adhesion and protease activities. They are involved in highly diverse biological processes, including spermatogenesis, neurogenesis, myogenesis, embryonic TGF-α release, and the inflammatory response(Edwards *et al*., 2008; Primakoff and Myles, 2000). To date, however, the function of most ADAMs in the brain is unknown. ADAM10 is widely expressed in different organs, including the brain. Over 40 substrates have been reported for ADAM10, including Notch, APP, PD-L1, Delta-like ligand-1 (Dll1), CX3CL1/Fractalkine, and N-cadherin(Edwards *et al*., 2008; Smith et al., 2020). ADAM10 regulates Notch signaling(Hsia et al., 2019; Kato et al., 2018; Lambrecht et al., 2018). It is implicated in neurological developmental disorders such as MECP2 duplication syndrome (MDS)(Wang et al., 2019). ADAM10 is also the main α-secretase to cleave the amyloid precursor protein(Kuhn et al., 2010). As a result, it is closely involved in neuronal development and the development of Alzheimer’s disease(Edwards *et al*., 2008; Postina et al., 2004).

ADAM10 is vital for the development of the CNS. Whole-body *Adam10* deletion in mice causes severe developmental disorders. *Adam10* deficiency causes lethality at an early embryonic stage (embryonic day 9.5 [E9.5]) alongside severe defects in brain development(Hartmann et al., 2002). The function and mechanisms of *Adam10* in development, especially during brain development, are not fully understood. In mice with *Adam10* deletion in neural progenitor cells using *Nestin-Cre;Adam10^fl/fl^ (Nestin^Adam10-CKO^)*, newborn pups die perinatally with a disrupted neocortex and ganglionic eminence(Jorissen et al., 2010). Early depletion of neural progenitor cells (NPCs) was observed due to precocious neuronal differentiation(Jorissen *et al*., 2010). To date, it is poorly understood whether ADAM10 regulates brain development by participating in gliogenesis. To answer this question, we performed genetic removal of *Adam10* from glial progenitors in transgenic mouse line *hGFAP-Cre;Adam10^fl/fl^* to study the role of ADAM10 in glial generation and development in the brain.

## Results

### Locomotor defects in mice after deletion of *Adam10* from glial progenitors

To investigate the role of ADAM10 in glial development, we performed conditional knockout of *Adam10 (Adam10-CKO)* in glial progenitor cells (GPCs) using *hGFAP-Cre;Adam10^fl/fl^* mice (hereafter, *hGFAP^Adam10-CKO^*). We observed that these mice had less fur than controls (i.e., wild type, *hGFAP-Cre*, *Adam^fl/+^*, and *Adam^fl/fl^* mice) beginning at postnatal day 8 (P8) (Figure 1A). They were nearly nude at P30 (Figure 1A). Compared with the control mice, the body weight of *hGFAP^Adam10-CKO^* mice started dropping at P5 and was significantly lower than that of the control group at P17 (Figure 1B). *hGFAP^Adam10-CKO^* mice survived to weaning (>P21), but all mice died before 4 months of age (n = 14 of 14; Figure 1C), which is very different from *Nestin^Adam10-CKO^* mice with *Adam10* deletion in neuronal stem cells; these mice die perinatally or shortly after birth(Jorissen *et al*., 2010). Because most glia are generated postnatally(Bandeira et al., 2009; Freeman, 2010; Ge et al., 2012), further analysis of glial development with *Nestin^Adam10-CKO^* is not feasible(Jorissen *et al*., 2010).

**Figure 1.**
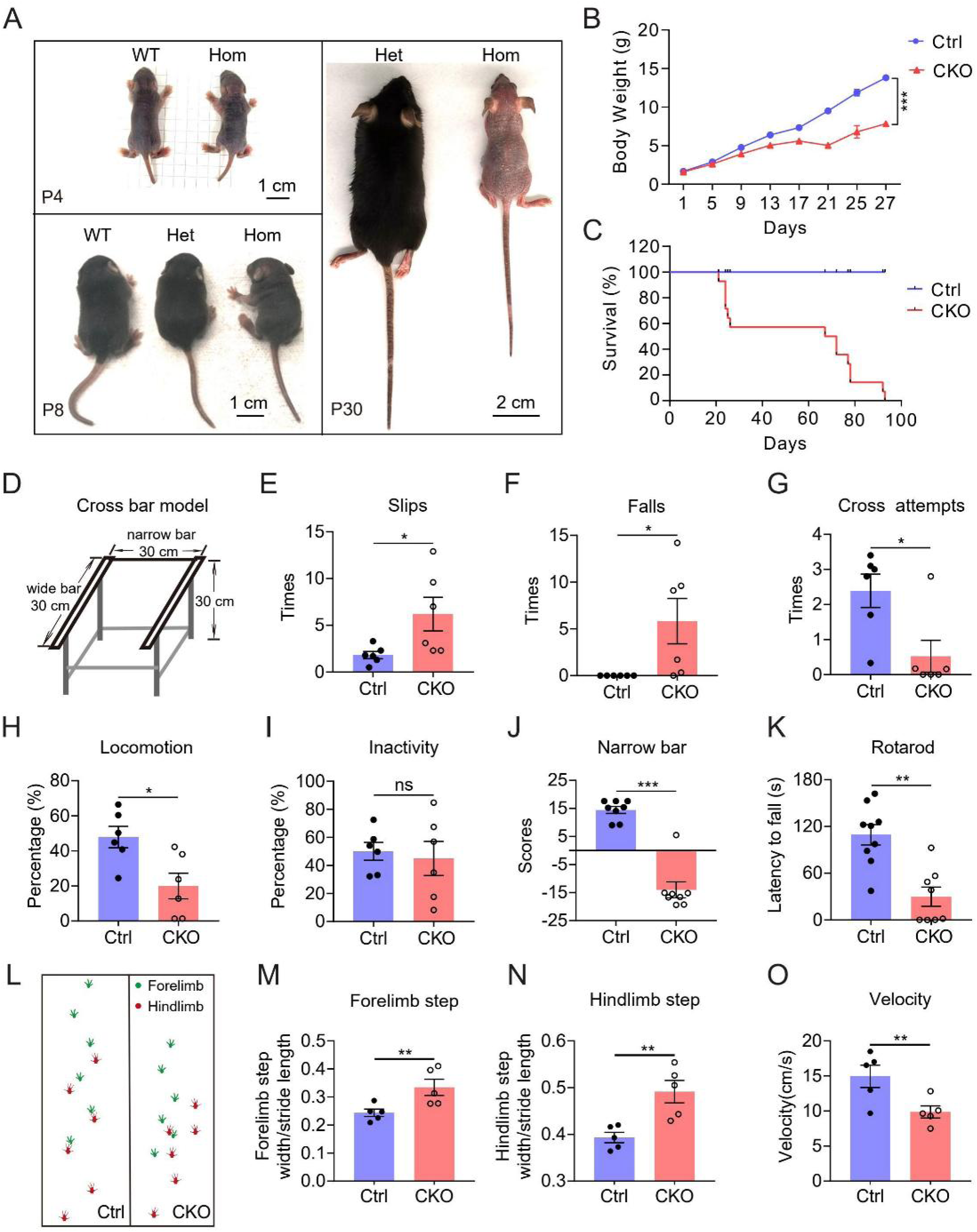
Phenotype of hGFAP-Cre;Adam10^fl/fl^ (hGFAP^Adam10-CKO^) mice. (**A**) Images of wild-type (WT, left), *hGFAP^Adam10-CKO^* (Hom, *hGFAP-Cre;Adam10^fl/fl^,* right), and heterozygous (Het, *hGFAP-Cre;Adam10^fl/+^,* middle) mice at P4, P8, and P30. (**B**) Body weight of *hGFAP^Adam10-CKO^* (CKO) and control (Ctrl) mice from P1 to P27. (**C**) Overall survival of both CKO and control mice. (**D**) A schematic of the bar-crossing apparatus for behavioral tests. (**E**) Frequency of slips from the wide bar. (**F**) Frequency of falls from the wide bar. (**G**) Frequency of attempts to cross from one wide bar to the other via the narrow bar. (**H**) Percentage of time spent in motion on the wide bar. (**I**) Percentage of time spent in inactivity on the wide bar. (**J**) A forced crossing score on the narrow bar was calculated for control and CKO mice. (**K**) The latency until falling for control and CKO mice during the rotarod test. (**L–O**) Graphical representation of selected gait parameters of control and CKO mice. Forelimb and hindlimb steps were recorded for analysis (L). Green, forelimb; red, hindlimb. The length of forelimb steps (M), the length of hindlimb steps (N), and the velocity (O) were then determined. The analyses were based on data shown in panel L. Mouse numbers: n = 11, 17, 17, 17, 17, 17, 7, and 2 mice for each time point of the control group, and n = 8, 14, 14, 14, 14, 13, 5, and 2 mice for the CKO group in B; n = 6 for both wild type and CKO in E–I; n = 14 for both CKO and control mice; n = 8 for wild type and CKO in J; n = 9 for wild type and 8 for CKO in K; and n = 5 for both wild type and CKO in M–O. Two-tailed unpaired *t*-test: \**p* < 0.05, \*\**p* < 0.01, \*\*\**p* < 0.005; ns, not significant. Error bars, s.e.m.

We performed multiple behavioral tests on balance and locomotion in *hGFAP^Adam10-CKO^* mice shortly after they were weaned (P21–30). A bar-crossing apparatus was used for balance and motor coordination tests (Figure 1D). With this apparatus, we observed significant differences in the slips, falls, crossing attempts, locomotion, inactivity, forced narrow bar crossings, and rotarod tests between *hGFAP^Adam10-CKO^* and control mice (Figure 1E–K). *hGFAP^Adam10-CKO^* mice slipped and fell from the bar with higher frequencies and made fewer attempts to cross the narrower bar (Figure 1E–G). The duration of locomotion on the wide bar was shorter, although the immobility duration on the wide bar was not different between control and *hGFAP^Adam10-CKO^* mice (Figure 1H, I). The scores for forced crossing tests on the narrow bar in *hGFAP^Adam10-CKO^* mice were significantly lower than those for the mice in the control group (Figure 1J). In the rotarod tests, *hGFAP^Adam10-CKO^* mice fell within a shorter period than the control group (Figure 1K). *hGFAP^Adam10-CKO^* mice showed more severe gait abnormalities and balance problems (Figure 1L). There were also more substantial defects in their forelimb step width, hindlimb step width, and the velocity of their walking behavior (Figure 1L–O). In short, the deletion of *Adam10* in *hGFAP^Adam10-CKO^* mice leads to defects in their body weight, survival, and locomotor activities.

### Abnormalities of cerebellar development in *hGFAP^Adam10-CKO^* mice

To further elucidate the cellular mechanism underlying these behavioral deficits, we dissected the brains of *hGFAP^Adam10-CKO^* mice at different developmental stages. We did not observe a noticeable abnormality in the morphology of P0 brains from *hGFAP^Adam10-CKO^* mice, except there was bleeding in the cortical region close to the midline. Interestingly, the bleeding phenotype ameliorated at P5 and gradually disappeared by P6–10 (Figure 2A).

**Figure 2.**
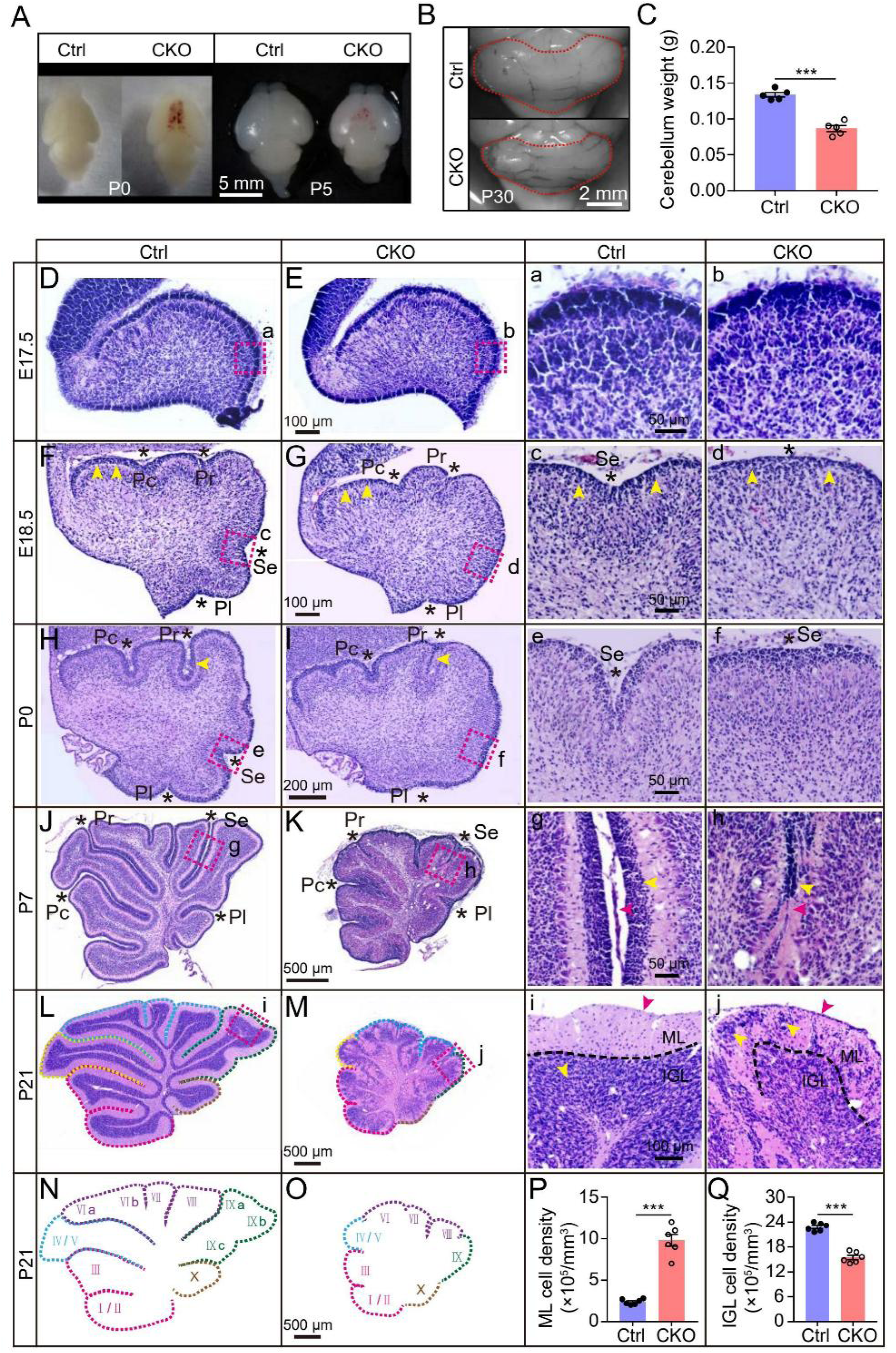
Cerebellar phenotype of *hGFAP^Adam10-CKO^* mice. (**A**) Images of the brains of *hGFAP^Adam10-CKO^* (CKO) and wild-type control (WT) mice at P0 and P5. (**B**) Images of the cerebellums of CKO and WT mice at P30. (**C**) Cerebellar weight of CKO (n = 5) and control (n = 5) mice at P30. (**D**, **E**) Hematoxylin and eosin (HE) staining of cerebellar sections from E17.5 control (D) and CKO **(E)** embryos. (a, b) Higher magnification of the boxed areas in D and E, respectively. (**F**, **G**) HE staining of cerebellar sections of E18.5 control and CKO embryos. Pc, Pr, Se, Pl (asterisks), the four primary fissures in the cerebellum. (c, d) Higher magnification of the boxed areas in F and G, respectively. Yellow arrowheads, EGL layers. (**H**, **I**) HE staining of P0 cerebellar sections of control and CKO mice. Yellow arrowheads, anterodorsal lobe and central lobe on either side of the primary fissure. (e, f) Higher magnification of the boxed areas in H and I, respectively. (**J**, **K**) HE staining of cerebellar sections of P7 control and CKO mice. (g, h) Higher magnification of the boxed areas in J and K, respectively. Lobes around Pr, Se, and Pl were attached in CKO mice (K, h: red arrowheads). Granule cell progenitors in EGL layers are shown (g, h: yellow arrowheads). (**L**, **M**) HE staining of cerebellar sections of P21 control and CKO mice. (i, j) Higher magnification of the boxed areas in L and M, respectively. Disorganized lamination of the cerebellar cortex in CKO mice (j). EGL,external granule layer (red arrowhead), ML, molecular layer; IGL, internal granule layer. Granule cell progenitors remained in the EGL in CKO mice (j, red arrowhead) but not in control mice (i, red arrowhead). Granule cell progenitor also stuck in the ML in CKO mice (j, yellow arrowhead) but migrated to IGL in control mice (i, yellow arrowhead). Dashed line in i and j, the border between the ML and IGL. (**N**, **O**) Representative diagram of the sagittal plane of the cerebellum of P21 control (N) and CKO (O) mice. (**P**, **Q**) Comparison of the cell density in the ML (P) and IGL (Q) layers from P21 cerebellums of control and CKO mice. Two-tailed unpaired *t*-test: \*\*\**p* < 0.001. Error bars, s.e.m.

The most apparent difference in the brains of *hGFAP^Adam10-CKO^* mice occurred in the cerebellum, which was significantly smaller in *hGFAP^Adam10-CKO^* mice relative to control mice (Figure 2B, C). After performing hematoxylin and eosin (HE) staining, we observed that the cerebellums of both *hGFAP^Adam10-CKO^* and control mice showed a curved, sausage-like structure with a smooth surface and no sulcus cleft formation at E17.5 (Figure 2D). No significant difference between *hGFAP^Adam10-CKO^* and control mice was apparent in the cerebellar cortical cells (Figure 2D, E). At E18.5, fissure Se was absent in the cerebellum of *hGFAP^Adam10-CKO^* mice, and the cell density in the external granule layer (EGL) of the *hGFAP^Adam10-CKO^* cerebellum was lower than in control mice (Figure 2F, G). At P0, fissure Se began to form in the cerebellum of *hGFAP^Adam10-CKO^* mice, although the folding in fissure Se in the control mice were more pronounced than in *hGFAP^Adam10-CKO^* mice (Figure 2H, I). Granule cell precursors (GCPs) in *hGFAP^Adam10-CKO^* mice accumulated in fissure Se (Figure 2I). The region of fissure Pl in *hGFAP^Adam10-CKO^* mice was shallower than in control mice (Figure 2H, I). In addition, the anterodorsal lobe and the central lobe around the primary fissure in *hGFAP^Adam10-CKO^* mice were attached, whereas they were separated in control mice (Figure 2H, I).

At P7, the size of the cerebellum of *hGFAP^Adam10-CKO^* mice relative to control mice was smaller (Figure 2J, K). The four primary fissures (Pc, Pr, Se, and Pl) had developed by this time point in both control and *hGFAP^Adam10-CKO^* mice (Figure 2J, K). Cortical lamination in the cerebellum of *hGFAP^Adam10-CKO^* mice was, however, disorganized, and the lobes around Pr, Se, and Pl were attached (Figure 2K). Cerebellar meninges in *hGFAP^Adam10-CKO^* mice were present along the outer surface of the cerebellar cortex, whereas meninges extended deeply into fissures between neighboring lobes in control mice (Figure 2J, K). There was a thick EGL layer in control mice but a much thinner one in *hGFAP^Adam10-CKO^* mice. GCPs gathered in clusters in *hGFAP^Adam10-CKO^* mice (Figure 2J, K).

At P21, the *hGFAP^Adam10-CKO^* cerebellum was significantly smaller than that of control mice (Figure 2L, M), and organized lamination of the cerebellar cortex was lost in *hGFAP^Adam10-CKO^* mice. Some GCPs in *hGFAP^Adam10-CKO^* mice were stuck in the molecular layer (ML), and some remained along the pial surface (Figure 2M). The border between the ML and the internal granule layer (IGL) was unclear in *hGFAP^Adam10-CKO^* mice. Although the total number of folia in *hGFAP^Adam10-CKO^* mice was equal to that of the control mice, each folium in the knockout mice had a much smaller size, disorganized lamination, and unsegregated fissures. The cell density in the *hGFAP^Adam10-CKO^* ML was significantly higher than that in the control (Figure 2P), but the cell density of the IGL in *hGFAP^Adam10-CKO^* mice was about half that in the control (Figure 2Q). These results indicated that CKO of *Adam10* indeed causes developmental abnormalities of the cerebellum. Because motor behaviors, including gait and balance coordination, largely depend on cerebellar function, cerebellar morphological defects may explain the behavioral abnormalities in *hGFAP^Adam10-CKO^* mice.

Given that most astrocytes and oligodendrocytes are generated during postnatal stages(Canoll and Goldman, 2008; Ge *et al*., 2012), we chose the promoter of human GFAP to drive expression of Cre recombinase in the hGFAP-Cre mice, as it is active during the late embryonic stage (E15.5). Its expression was limited to specific forebrain regions, including the cerebral cortex and the subventricular zone (SVZ) and VZ regions (Figure S1). This result is consistent with the fate-mapping of tdTomato-expressing cells from *hGFAP-Cre;Ai14* mice. We observed that astrocytes, granular cells, and oligodendrocytes were labeled by red fluorescent protein in the cerebellum, but we did not observe that tdTomato was expressed in Purkinje cells (Figure S2). In short, *hGFAP^Adam10-CKO^* mice provided a valuable tool for studying the role of ADAM10 in early gliogenesis. Compared with *hGFAP-Cre*, the expression of Cre recombinase occurred at a much later stage in *mGFAP-Cre;Ai14* mice. We found that the tdTomato-expressing cells in the cerebral cortex and cerebellum of P2 mouse brains were mainly limited to astrocytes. This result strongly indicated that Cre recombinase expression appears after the determination of the astrocyte fate decision. It might explain why we could not detect defects in the cerebellum in *mGFAP-Cre;Adam10^fl/fl^* (*mGFAP-Cre^Adam10-CKO^*) mice (data not shown).

### *Adam10* deletion in GPCs alters the pathways for macroglial generation

To explore the underlying molecular mechanisms of cerebellar defects of *hGFAP^Adam10-CKO^* mice, we performed bulk RNA sequencing (bRNA-seq) with whole cerebellums of P0, P7 *hGFAP^Adam10-CKO^* and control mice (*hGFAP-Cre, Adam10^fl/+^,* and wild types, control data from the three groups were pooled; Figure 3A). In an unsupervised principal component analysis (PCA), samples from P7 *hGFAP^Adam10-CKO^* and control mice were well separated into two clusters. However, there were overlaps between P0 control and *hGFAP^Adam10-CKO^* samples (Figure 3B–G), which is consistent with the morphological results at different developmental stages: the morphological alteration in the cerebellums of *hGFAP^Adam10-CKO^* mice was minor in perinatal pups (i.e., E18.5 and P0; Figure 2F–I), but the size and folding features changed dramatically at P7 (Figure 2J, K).

**Figure 3.**
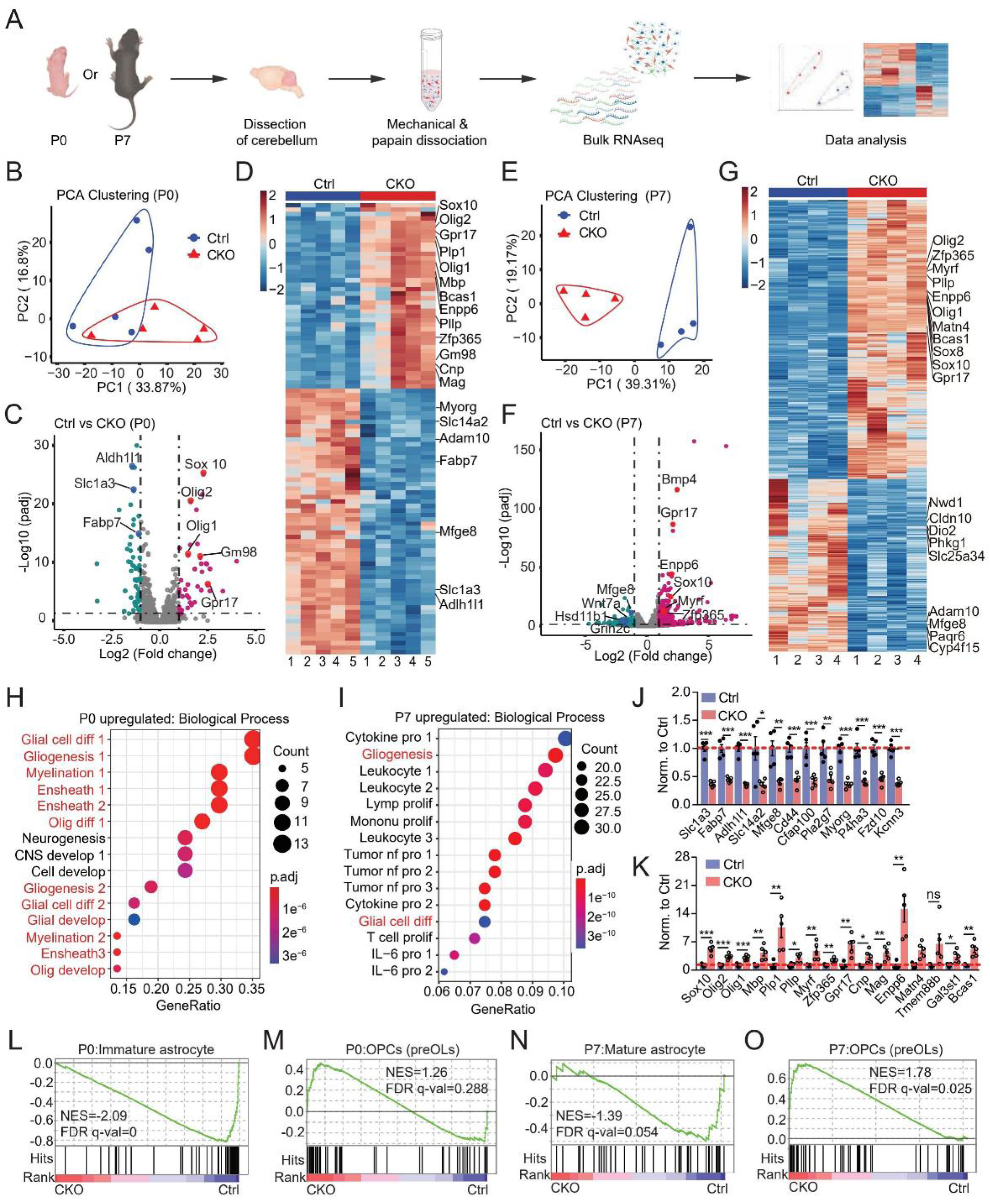
bRNA-seq analyses show that *Adam10* deletion altered pathways involved in glial generation and development. (**A**) Flow diagram of bRNA-seq of P0 and P7 cerebellums from control (Ctrl) and *hGFAP^Adam10-CKO^* (CKO) mice. (**B**–**F**) Principal component analysis (PCA) of bRNA-seq data from P0 (B) and P7 (E) cerebellar samples of control (n = 5, P0; n = 4, P7) and CKO (n = 5, P0; n = 4, P7) mice. (B) The top two PCs explain 33.87% and 16.8% of the total variance, respectively. (E) The top two PCs explain 39.31% and 19.17% of the total variance, respectively. Each circle indicates data from bRNA-seq of individual samples (C, F). The representative genes (e.g., Slc1a3, Olig2) are shown by a larger circle of a slightly different color. Analysis of differentially expressed genes (upregulated or downregulated) in each sample of control and *hGFAP^Adam10-CKO^* mice. (C, F) The volcano plots show downregulated (green and dark blue) and upregulated (dark pink and red) genes (FDR adjusted *p* < 0.05 and **|**log_2_FC**|** > 1); black vertical dashed lines highlight a FC of –1.5 and 1.5, and the black horizontal dashed line represents an adjusted *p*-value (*P*adj) of 0.05. (D, G) Clustered heatmaps for P0 mice (D) (control, n = 5; CKO, n = 5) and P7 mice (G) (control, n = 4; CKO, n = 4). The ordinate represents individual genes, and the abscissa represents individual samples. The colors of the heatmap correspond to the expression level of genes (D, G). Red indicates high expression and blue indicates low expression. DEGs are grouped by hierarchical clustering analysis, based on Euclidean distance**s** and the ward.D2 method. (**H, I**) Gene Ontology (GO) biological process enrichment analysis of upregulated pathways in P0 (H) and P7 (I) mice. The process names highlighted in red are related to glial generation and development: Glial cell diff 1, glial cell differentiation; Gliogenesis 1, gliogenesis; Myelination 1, myelination; Ensheath 1, ensheathment of neurons; Ensheath 2, axon ensheathment; Oligo diff 1, oligodendrocyte differentiation; Neurogenesis, negative regulation of neurogenesis; CNS develop 1, negative regulation of nervous system development; Cell develop, negative regulation of cell development; Gliogenesis 2, regulation of gliogenesis; Glial cell diff 2, regulation of glial cell differentiation; Glial develop, glial cell development; Myelination 2, central nervous system myelination; Ensheath 3, axon ensheathment in the central nervous system; Olig develop, oligodendrocyte development; Cytokine pro 1, positive regulation of cytokine production; Leukocyte 1, leukocyte**-**mediated immunity; Leukocyte 2, leukocyte proliferation; Lymp prolif, lymphocyte proliferation; Mononu prolif, mononuclear cell proliferation; Leukocyte 3, myeloid leukocyte activation; Tumor nf pro 1, tumor necrosis factor production; Tumor nf pro 2, tumor necrosis factor superfamily cytokine production; Tumor nf pro 3, regulation of tumor necrosis factor production; Cytokine pro 2, regulation of tumor necrosis factor superfamily cytokine production; Glial cell diff, glial cell differentiation; T cell prolif, T**-**cell proliferation; IL-6 pro 1, interleukin-6 production; IL-6 pro 2, regulation of inteleukin-6 production. (**J**, **K**) Relative abundance of astrocyte- (J) and oligodendrocyte- (K) specific genes from P0 control (blue, n = 5 mice) and CKO (red, n = 5 mice) samples. Each circle represents individual sample. Two-tailed unpaired *t*-test: \**p* < 0.05, \*\**p* < 0.01, \*\*\**p* < 0.005. Error bars, s.e.m. (**L–O**) The Gene Set terms immature astrocyte**s** (L), OPCs (pre-OLs, i.e., oligodendrocyte progenitor cells and pre-myelinating oligodendrocytes) (M, O), and mature astrocytes (N) were differentially regulated in cerebellar samples from P0 and P7 CKO mice when compared with the control samples in a GSEA analysis. Vertical black bars indicate the hits in the Gene Sets as represented among all genes pre-ranked by ranking metrics. The green dashed lines in the plots indicate thresholds of *p*adj < 0.05 and |Log_2_(FC)| > 1.

Compared with the control samples, there were 60 downregulated genes and 42 upregulated genes in P0 *hGFAP^Adam10-CKO^* samples (fold change (FC) > 2, *p* < 0.05; Figure 3D). Interestingly, we noticed that most of the downregulated genes were astrocyte-enriched genes, including *Aldh1l1*, *Slc1a3*, *Myorg*, *Slc14a2*, and *Fabp7*. However, the upregulated genes were mainly oligodendrocyte lineage–enriched genes, including *Cnp*, *Plp1*, *Bcas1*, *Enpp6*, *Mbp*, and *Mag*, which are highly expressed in newly formed and mature oligodendrocytes(Zhang et al., 2014). *Olig1*, *Sox10*, *Olig2*, *Pllp*, and *Zfp365,* which are highly expressed in the oligodendrocyte lineage (i.e., oligodendrocyte progenitor cells [OPCs] and newly formed oligodendrocytes), were also found on the list (Figure 3C, D).

Similar results indicating upregulation of oligodendrocyte-enriched genes were also detected in P7 *hGFAP ^Adam10-CKO^* samples (Figure 3E–G), although at this stage we detected more differentially expressed genes between *hGFAP^Adam10-CKO^* and control samples, including *Bmp4*, *Gpr17*, *Enpp6*, *Sox10*, *Myrf*, and *Zfp365*. Gene Ontology (GO) analysis of P0 bRNA-seq results revealed that most of these upregulated genes belonged to the gene sets Glial Cell Differentiation, Gliogenesis, Myelination, Ensheath, and Oligodendrocyte Development. GO analysis of the P0 and P7 results indicated that Gliogenesis and Glial Cell Differentiation were among the top gene sets altered in the transcriptomes of the cerebellums from *hGFAP^Adam10-CKO^* mice (Figure 3H, I). In addition, the mRNA abundance of astrocyte-specific genes was significantly decreased in *hGFAP^Adam10-CKO^* cerebellums (Figure 3J, S3A), but oligodendrocyte-specific genes increased (Figure 3K, S3B).

Using pathway analysis with Gene Set Enrichment Analysis (GSEA), we observed that astrocyte-associated pathways were more active in the control P0 and P7 samples (Figure 3L, N). However, OPC- or pre-mature oligodendrocyte (pre-OL)–associated pathways were more active in *hGFAP^Adam10-CKO^* mice of the same ages (Figure 3M, O). With our analysis of bRNA-seq data from the cerebellum of *hGFAP^Adam10-CKO^* mice, we saw similar changes to the transcriptomes pertaining to glial cell development and gliogenesis. The pathways related to astrocyte generation and development were inhibited, whereas the pathways related to oligodendrocyte generation and development were significantly enhanced in *hGFAP^Adam10-CKO^* mice. Our results strongly suggested that ADAM10 plays an essential role in the generation of astrocytes and oligodendrocytes in the developing brain.

### *Adam10* deletion inhibits astrocyte generation but promotes oligodendrocyte production

To further investigate whether the deletion of *Adam10* in GPCs alters the generation of astrocytes and oligodendrocytes, we collected cerebellums from P2 *hGFAP^Adam10-CKO^* and wild-type mice and prepared cell suspensions for single-cell RNA-seq (RNA-seq) analysis (Figure 4A). To analyze the fate of GPCs and determine their progeny, we used *hGFAP-Cre;Adam10;Ai14* and *hGFAP-Cre;Ai14* (control) mice for measurements. A red fluorescent protein, tdTomato, was expressed in GPCs and their progeny once Cre recombinase removed the stop codon in the *Ai14* reporter line. Among all pooled cells for single-cell transcriptome analysis from both *hGFAP^Adam10-CKO^* and control cerebellums, we obtained 1,164 tdTomato-positive cells for further analysis. To classify the major cell types in the neonatal cerebellum, we performed *t*-distributed stochastic neighbor embedding (*t-*SNE) analyses [<u>9</u>]. As a result, we identified 14 major cell clusters from the developing cerebellum: astrocytes, cells from the choroid plexus, endothelial cells, excitatory neurons, GABAergic cells, granule cells, interneurons, microglia, oligodendrocytes, Purkinje cells, unipolar brush cells, and three other neuron clusters (Figure 4B). Every cell cluster was determined by two or more specific gene markers (Figure 4D). For example, we identified the astrocyte cluster with *Aldh1l1* and *Pla2g7* and the oligodendrocyte cluster with *Sox10* and *Olig2*. We noticed that the distribution pattern of different types of cells differed between *hGFAP^Adam10-CKO^* and control samples (Figure 4C).

**Figure 4.**
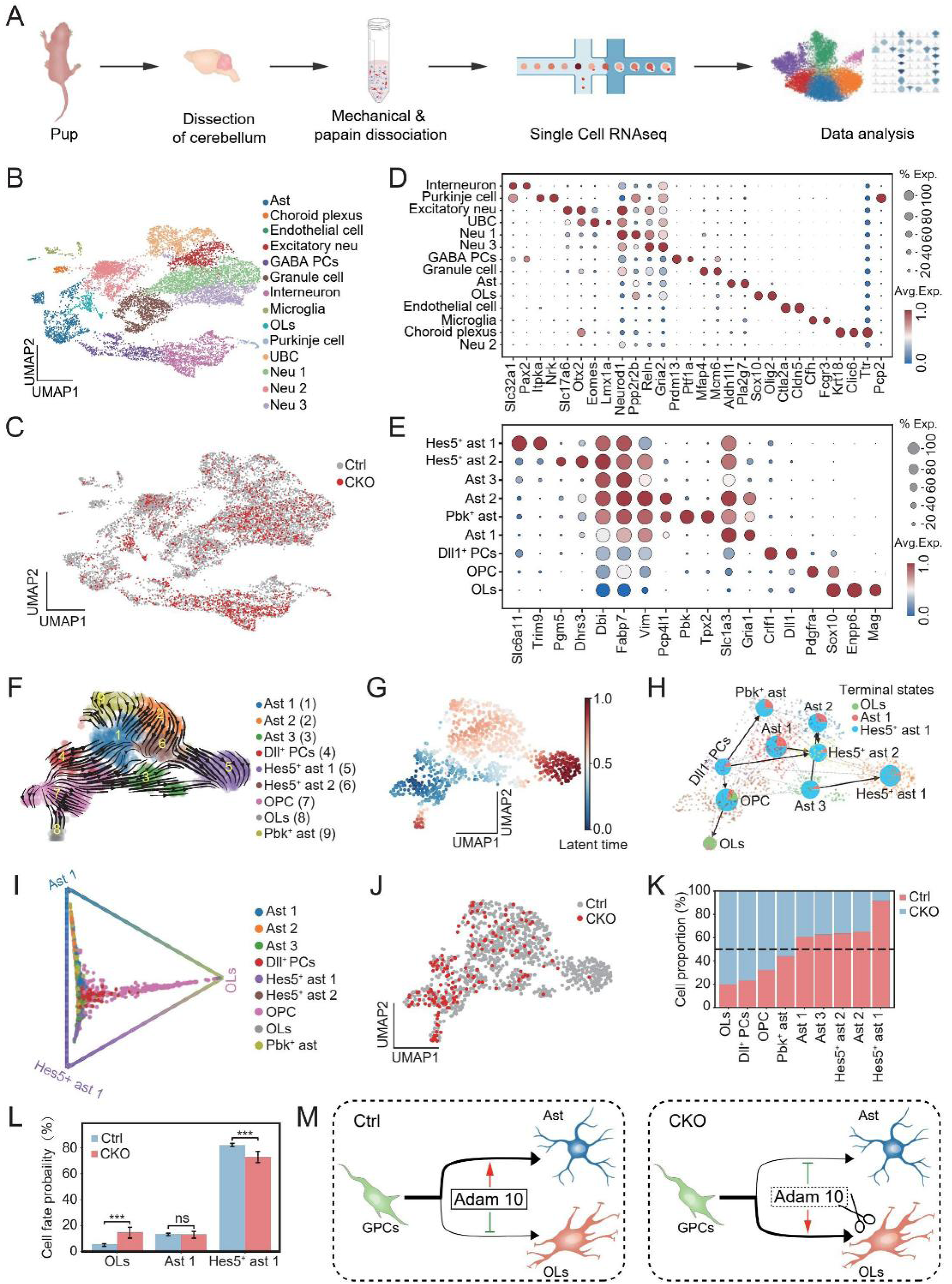
The absence of *Adam10* inhibits astrocyte generation but promotes oligodendrocyte generation in the cerebellum. (**A**) Schematic diagram of analysis via single-cell RNA-seq from P2 mouse cerebellums of *hGFAP-Cre;Ai14* (control, Ctrl) and *hGFAP-Cre;Adam10^fl/fl^;Ai14* (CKO) mice. (**B**) Visualization of major classes of cells using nonlinear dimensionality reduction technique, uniform manifold approximation and projection (UMAP). Each dot represents an individual cell; different colors represent different cell classes. Neu 1–3, neurons; PCs, progenitor cells; Ast, astrocytes; OLs, oligodendrocytes; UBC, unipolar brush cells, Excitatory neu, excitatory neurons; GABA PCs, GABAergic progenitor cells. The total number of cells analyzed was 11,155 from CKO and control mice at P2. (**C**) Distribution of the 14 identified subclusters of cells in the cerebellums of control and CKO mice. Red, cells with tdTomato expression from CKO mice. (**D**) Dot plots of marker genes for the corresponding subclusters listed in (B). (**E**) Dot plots of marker genes for glial progenitors, astrocytes, and oligodendrocytes labeled by tdTomato in the cerebellums of control and CKO mice. PCs, progenitor cells; Ast, astrocytes; OLs, oligodendrocytes; OPC, oligodendrocyte precursor cells. (**F**) UMAP of astrocytes and oligodendrocytes from P2 control and CKO cerebellums. Streamlines show averaged and projected scVelo velocities. Each cluster type is represented by a number in the stremlines. The glial cell subclusters include six astrocyte clusters (Ast1-3, Hes^+^ ast1, Hes^+^ ast2, and Pbk^+^ ast), two oligodendrocyte cell clusters (OPC, OLs), and Dll^+^ progenitor cells (Dll^+^ PCs). (**G**) CellRank probabilities for acquiring the terminal cell fate. Cells were colored based on the probability of reaching the differentiated state. Dark red and blue show the most differentiated and undifferentiated states, respectively. This latent time indicates internal clock of a cell and approximates the real time experienced by a cell as it differentiates. (**H**) Differentiation pathway connectivity of progenitor cells and eight different glial subclusters from the partition-based graph abstraction (PAGA) shown in (F). The proportion of the pie chart represents the probability of each terminal state, and the arrow represents the direction of cell differentiation. (**I**) Triangle projection of 1,164 tdTomato-expressing cells according to fate probabilities. Each color represents a glial subcluster. Macrostates are arranged on the edge of a triangle; each cell is placed inside the triangle according to its probability of reaching any terminal state. Cells in the center have a higher multilineage potential, whereas cells close to one of the corners are committed to differentiate into that terminal cell type. The cell types consist of astrocytes (Ast1 and Hes5^+^ ast1) and oligodendrocytes (OLs). (**J**) Distribution pattern of cells from the cerebellum of the CKO (red) and control (gray) mice. (**K**) Proportions of cell subpopulations of astrocytes and oligodendrocytes in control and CKO cerebellums. (**L**) Cell fate probability of differentiation into oligodendrocytes (OLs) and astrocytes (Ast 1 and Hes5^+^ ast 1) in control and CKO mice. Two-tailed unpaired *t*-test: \*\*\**p* < 0.001; ns, not significant. Error bars, s.e.m. (**M**) Schematic diagram showing that *Adam10* mediates the generation of oligodendrocytes (OLs) and astrocytes (Ast) from glial progenitor cells (GPCs). ADAM10 promotes astrocyte generation under physiological conditions. Once *Adam10* is removed (as indicated by the scissors), the path for oligodendrocyte generation from GPCs is activated.

To further analyze the alterations in macroglial cell clusters and their related molecular features, we segregated the macroglial subcluster (including clusters of astrocytes and oligodendrocytes) into nine distinct glial subclusters with a random forest6 analysis and identified the top one or two genes in each subcluster (Figure 4E). We observed that Dll^+^ progenitor cells (Dll^+^ PCs) were located at the starting point and then differentiated into astrocytes and OPCs. OPCs further differentiated into oligodendrocytes (Figure 4F). According to the analysis of differentiation degree, Dll^+^ PCs were undifferentiated glial subclusters, whereas Hes5^+^ astrocytes (i.e., Hes5^+^ ast 1 in Figure 4H) and oligodendrocytes (i.e., OLs in Figure 4H) were the most differentiated subclusters (Figure 4G and 4H). Nearly all subclusters eventually differentiated into Hes5^+^ astrocytes (Figure 4H). Moreover, our results from fate probability analysis of the tdTomato-expressing cells in the cerebellum of *hGFAP-Cre;Ai14* mice suggested that Dll^+^ PCs differentiated into astrocytes (i.e., subclusters Ast 1 and Hes5^+^ ast 1) and OLs (Figure 4I). Simultaneously, we detected a difference in the relative cell numbers of these nine glial subclusters between *hGFAP^Adam10-CKO^* and control mice (Figure 4J). We observed that the proportions of OLs, Dll^+^ PCs, and OPCs increased substantially in *hGFAP^Adam10-CKO^* mice, whereas the proportions of astrocyte subclusters (Ast 1, Ast 2, Ast 3, Hes5^+^ ast 1, and Hes5^+^ ast 2) decreased (Figure 4K). Compared with the control group, the cell fate probability of differentiating into oligodendrocytes was higher in *hGFAP^Adam10-CKO^* mice, but the probability of differentiating into astrocytes was lower (Figure 4L). These comprehensive analyses of single-cell transcriptomes indicated that GPCs in the developing cerebellum tend to differentiate into astrocytes under physiological conditions, whereas they are more likely to differentiate into oligodendrocytes once *Adam10* is downregulated (Figure 4M). Thus ADAM10 functions as a master regulator for the generation of astrocytes and oligodendrocytes during cerebellar development.

### *Adam10* deletion promotes oligodendrocyte production in the cerebellum

To further verify whether the removal of *Adam10* in GPCs of *hGFAP^Adam10-CKO^* mice alters the macroglial cell fate, we performed immunostaining with various antibodies against astrocytes or oligodendrocytes in the cerebellums of control and *hGFAP^Adam10-CKO^* mice. Astrocytes were labeled with an antibody against brain lipid binding protein (Blbp, also known as Fabp7). We observed that the density of astrocytes (i.e., Blbp^+^ cells) decreased by ∼50% in the IGL and white matter (WM) of the cerebellums from P0 and P8 *hGFAP ^Adam10-CKO^* mice relative to control mice (Figure 5A–F). Further, we noticed that astrocytes were distributed in an orderly fashion near the Purkinje cell layer in P8 control mice, whereas they were disorganized and sparsely distributed in an irregular manner in the cerebellums of P8 *hGFAP^Adam10-CKO^* mice (Figure 5D–F), which is consistent with the HE staining results from P7 and P21 mice (Figure 2J–M).

**Figure 5.**
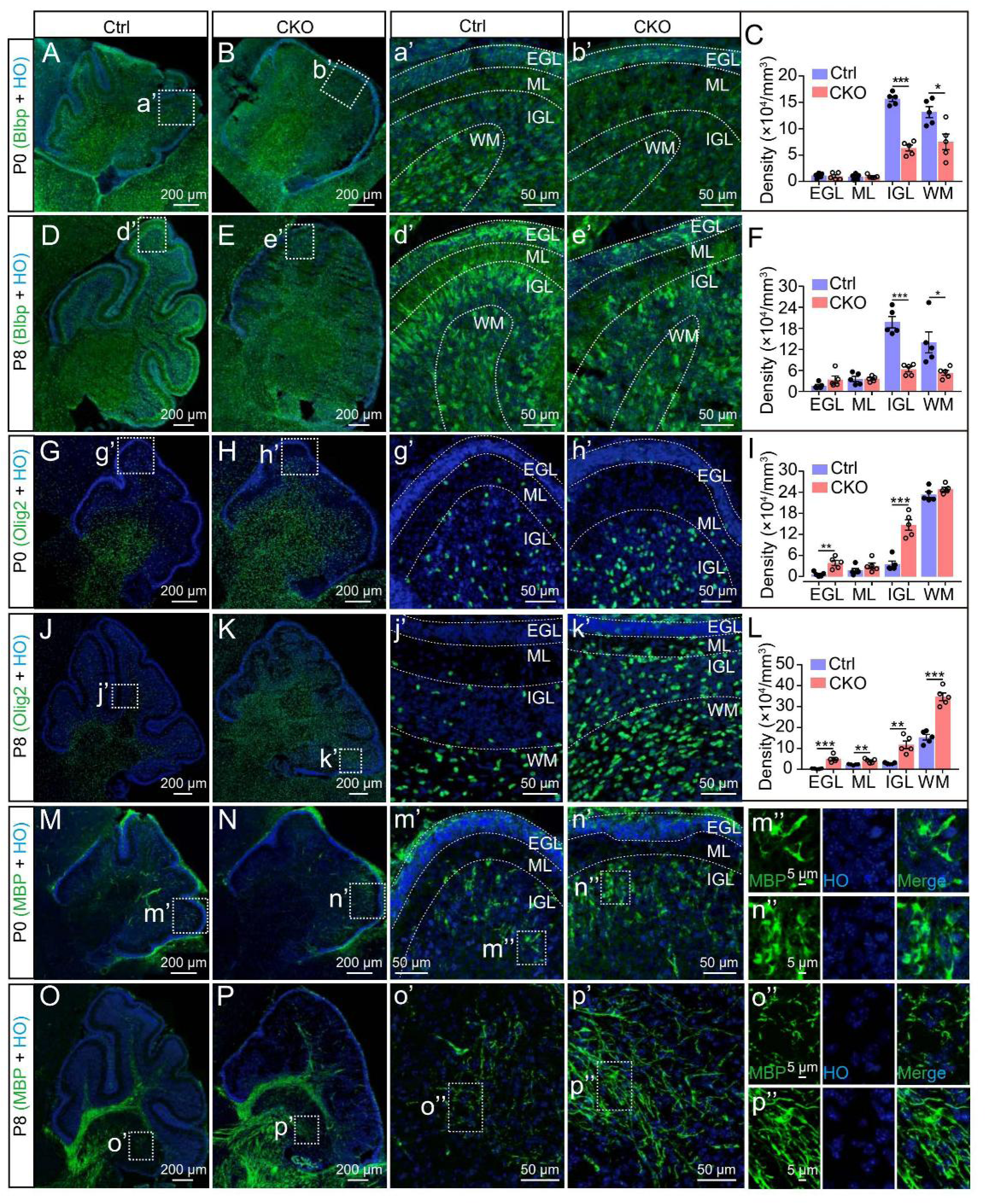
Staining of astrocytes and oligodendrocytes in the cerebellum after the removal of *Adam10* in glial progenitor cells. (**A–F**) Images of cerebellar sections and quantification from the cerebellum of wild-type control (Ctrl) and *hGFAP^Adam10-CKO^* (CKO) mice at P0 (A–C) and P8 (D–F). Green, cells stained with anti-Blbp; blue, nuclei stained with Hoechst 33342 (HO). Comparisons of the density of Blbp^+^ cells in different cerebellar subregions (EGL, ML, IGL, and WM) of control (A, D) and CKO (B, E) mice at P0 (A–C) and P8 (D–F). (a’, b’, d’, e’) Higher magnification of the boxed areas in A, B, D, E, respectively. Dashed lines indicate the margins of different layers in the cerebellum, including the external granule layer (EGL), molecular layer (ML), internal granule layer (IGL), and white matter (WM). The density (×10^4^ cells/mm^3^) of the IGL at P0: control, 15.66 ± 0.51; CKO, 6.3 ± 0.57; at P8: control, 19.75 ± 1.63; CKO, 6.30 ± 0.71. The density of the WM at P0: control, 13.14 ± 1.05; CKO, 7.51 ± 1.48; at P8: control, 13.00 ± 3.00; CKO, 5.31 ± 0.70. (**G–L**) Images of cerebellar sections from wild-type control and CKO mice at P0 (G–I) and P8 (J–L). Green, cells stained with anti-Olig2; blue, nuclei stained with Hoechst 33342 (HO). Comparisons of the density of Olig2^+^ cells in different cerebellar subregions (EGL, ML, IGL, and WM) of control (G, J) and CKO (H, K) mice. (g’, h’, j’, k’) Higher magnification of the boxed areas in G, H, J, K, respectively. (**M–P**) Images of cerebellar sections of wild-type control and CKO mice at P0 (M, N) and P8 (O, P). Green, myelination stained with anti-MBP; blue, nuclei stained with Hoechst 33342 (HO). Dashed lines indicate the margins of different layers in the cerebellum (EGL, ML, and IGL) of control (M, O) and CKO (N, P) mice at P0 (M, N) and P8 (O, P). (m’, n’, o’, p’) Higher magnification of the boxed areas in M, N, O, P, respectively. (m’’, n’’, o’’, p’’) Representative regions of MBP staining of oligodendrocyte myelination as indicated by the boxed areas in m’, n’, o’, p’, respectively.

To detect the distribution and maturation of the oligodendrocyte cell lineage in the cerebellum, we stained cerebellar sections with an antibody against Olig2. We observed that oligodendrocytes (Olig2^+^ cells) increased significantly in the EGL and IGL layers of P0 *hGFAP^Adam10-CKO^* mice. The apparent increase in oligodendrocyte density was also found in the EGL, ML, IGL, and WM layers of P8 *hGFAP^Adam10-CKO^* mice (Figure 5G–L). In addition, we used an antibody against MBP, a marker of mature oligodendrocytes, MBP expression was notably higher in the cerebellum of *hGFAP^Adam10-CKO^* mice, implying that more myelination occurred in the oligodendrocytes of *hGFAP^Adam10-CKO^* mice (Figure 5M–P). These results were consistent with our results from bRNA-seq: deletion of *Adam10* led to the upregulation of the myelination-related pathways (Figure 3H). These results demonstrate that the deletion of *Adam10* from GPCs promotes oligodendrocyte production and inhibits astrocyte generation in the developing cerebellum.

## Discussion

In this study, we investigated the contribution of ADAM10 to cerebellar development and observed that the conditional knockout of *Adam10* in GPCs of the developing mouse brain promoted the production of OPCs and oligodendrocytes but inhibited the generation of astrocytes. These results indicated that ADAM10 mediates gliogenesis in the developing brain.

ADAM10 cleaves the Notch receptor and promotes the expression of Notch-targeted genes, for example *Hes1/5*, *Hey1*, and *Dtxl(Hartmann et al., 2002; Kopan and Ilagan, 2009)*. The absence of ADAM10 would thus inhibit the activation of the Notch signaling pathway, which is involved in the regulation of neurogenesis and gliogenesis(Engler et al., 2018; Lasky and Wu, 2005; Wang and Barres, 2000). Notch signaling controls lateral inhibition and contributes to early neuronal differentiation(Lewis, 1996). Previous studies have shown that Notch activation inhibits neuronal differentiation and maintains neural progenitor cells (NPCs), but it promotes gliogenesis(Grandbarbe et al., 2003; Lütolf et al., 2002). During the early embryonic stage, Notch signaling is involved in the selection process of neural progenitors(Heitzler and Simpson, 1991). During the second trimester, neuroepithelial cells undergo asymmetric divisions. They generate neural-like intermediate neural progenitors (INPs) and radial glial cells. Neurons that differentiate from INPs express Notch ligands. Their ligands interact with the Notch receptors on nearby radial glial cells and activate the JAK/STAT pathway, promoting astrocyte differentiation(Namihira et al., 2009). The ligand Jag2 activates Notch receptors via the PMN domain (the progenitor domain of motor neurons) and prevents OPC generation by maintaining a high level of Hes5(Rabadán et al., 2012). From the late embryonic stage to P6 in mice, activation of the canonical Notch pathway by the ligand Jag1 induces the transcription of *Hes1* and prevents OPC maturation(Traiffort and Ferent, 2015). Loss of the transcription factor Hes1 leads to a reduction in the number of neural stem cells (NSCs) and an excessive increase in INPs(Del Bene et al., 2008; Imayoshi et al., 2013). In NSCs, downregulation of *Hes1* expression leads to *Ngn2* activation, triggering neuronal differentiation(Formosa-Jordan et al., 2013; Pierfelice et al., 2011). After P6, the non-canonical Notch pathway is activated by the ligand F3/contactin. This activation promotes oligodendrocyte generation from GPCs and leads to OPC differentiation(Cui et al., 2004; Hu et al., 2003).

Given the regulatory roles of the Notch pathway in neurogenesis and gliogenesis during brain development, we predicted that the potential phenotype in ADAM10 conditional knockout mice (*hGFAP^Adam10-CKO^*) might arise from dysregulation in the Notch pathway. We observed downregulated expression of Notch1 and Notch2 in P0 *hGFAP^Adam10-CKO^* mice (Figure S6). Therefore, gliogenesis, which requires the participation of Notch, is dysregulated due to the removal of *Adam10*. As shown in our results, the deletion of *Adam10* inhibited astrocyte generation, which is also consistent with the bRNA-seq results in which astrocyte-enriched genes were remarkably downregulated (Figure 3J). In addition, the removal of *Adam10* induced OPC generation and differentiation (Figures 3 and 5).

Our results reveal the effect of *Adam10* deficiency on gliogenesis in *hGFAP^Adam10-CKO^* mice. Neurogenesis in the mouse cerebral cortex starts at E12 and peaks at E14, whereas gliogenesis occurs from the late embryonic stage to the early postnatal period (E17–P4)(Sauvageot and Stiles, 2002). In the *hGFAP-Cre* mouse line, we observed that Cre expression started at E15.5, where it was expressed in multi-potential NSCs as previously reported(Zhuo et al., 2001). X-gal staining also showed that Cre expression was high in the SVZ/VZ of the neocortex at E15.5. Although there are NSCs in the SVZ/VZ of *hGFAP^Adam10-CKO^* mice, *Adam10* deletion primarily interferes with gliogenesis. In *Nestin-Cre* mice, the expression of Cre transcripts starts at E9.0 and occurs throughout the brain and spinal cord from E10.5(Yang et al., 2004). Cre is expressed in neural progenitor cells of the VZ(Graus-Porta et al., 2001). Compared with *Nestin^Adam10-CKO^* mice*, hGFAP^Adam10-CKO^* mice underwent *Adam10* deletion at much later stages (∼E13.5–15.5). Neurogenesis occurred at the expense of astrocytes in *Nestin^Adam10-CKO^* mice. Because neurogenesis precedes gliogenesis, it is hard to determine whether abnormalities in neurogenesis are followed by defects in gliogenesis in *hGFAP^Adam10-CKO^ mice.* Future studies of ADAM10 are needed to clarify this point.

## Supplementary Figure legends

**Supplementary figure 1.**
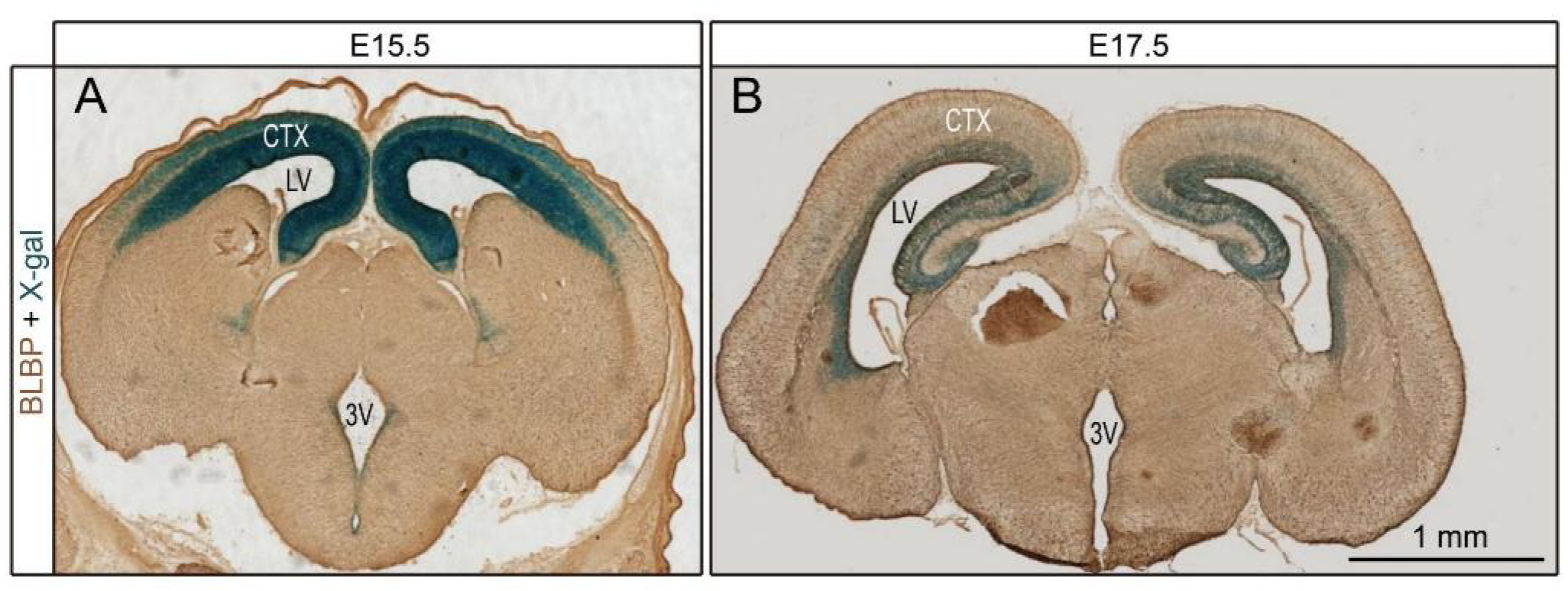
Expression of Cre in *hGFAP^Adam10-CKO^* mice. (**A**, **B**) The expression pattern of Cre recombinase in *hGFAP-Cre;Rosa26^Lacz^* embryos at E15.5 and E17.5. Blue, X-gal staining.

**Supplementary figure 2.**
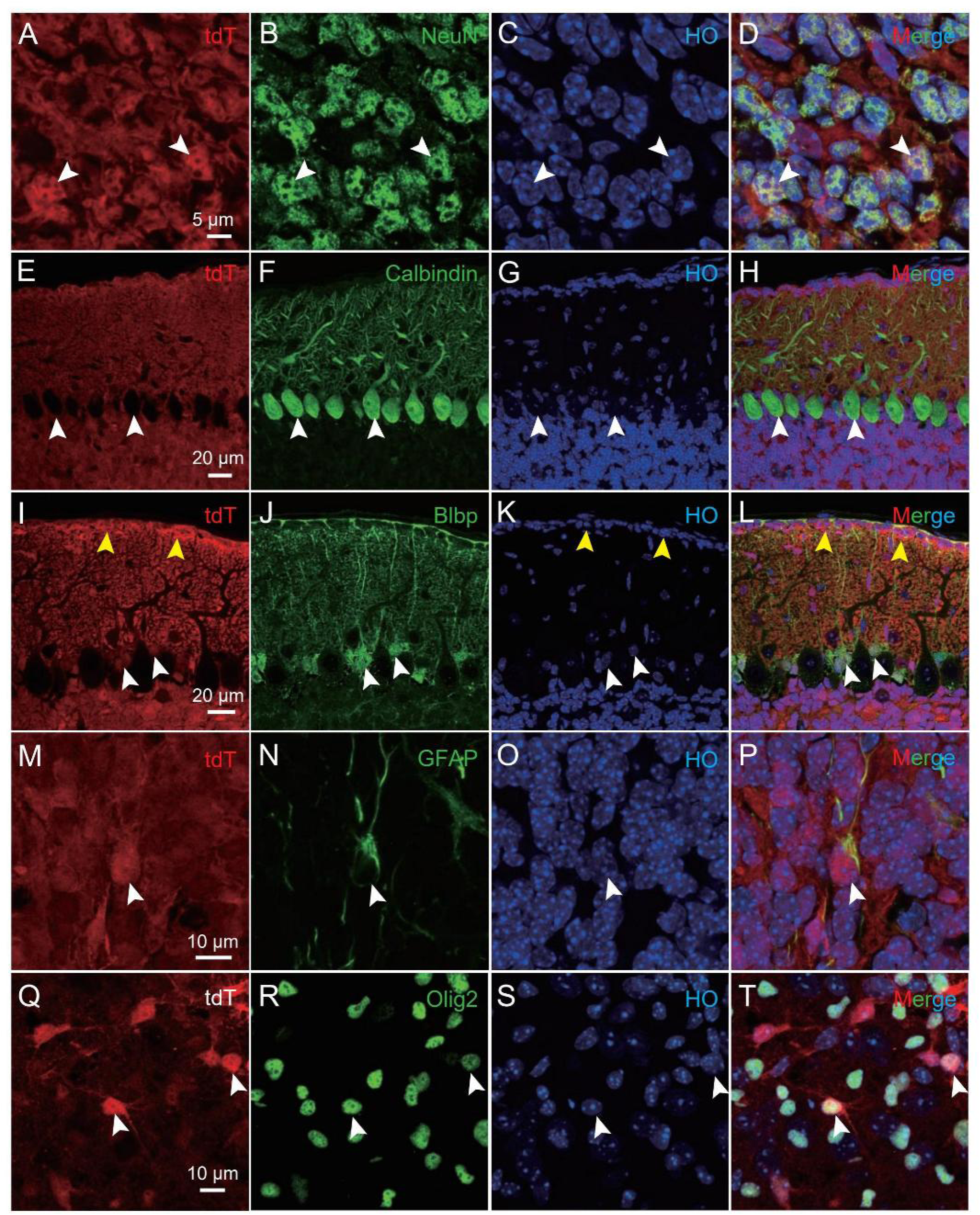
Characterization of tdTomato-expressing cells in the cerebellum of *hGFAP-Cre;Ai14* mice. (**A–T**) Images of cerebellar sections from *hGFAP-Cre;Ai14* mice at P14. Red, the fluorescent signals from tdTomato (Tdt; A, E, I, M, Q). Green, staining with anti-NeuN (B, neurons), anti-calbindin (F, Purkinje cells), anti-Blbp (J, astrocytes), anti-GFAP (N, astrocytes), and anti-Olig2 (R, oligodendrocytes and OPCs) to detect the indicated cell types. Blue, nuclei stained by Hoechst 33342 (C, G, K, O, S). Merged images shown in (D, H, L, P, T). White arrowheads in (A–T), tdTomato-expressing cells in (A), Purkinje cells in (E–H), immature astrocytes in (I–L), tdTomato-expressing cells in (M–P), and cells of the oligodendrocyte lineage in (Q–T); Yellow arrowheads in (I, K, L), glial endfeet in the EGL.

**Supplementary figure 3.**
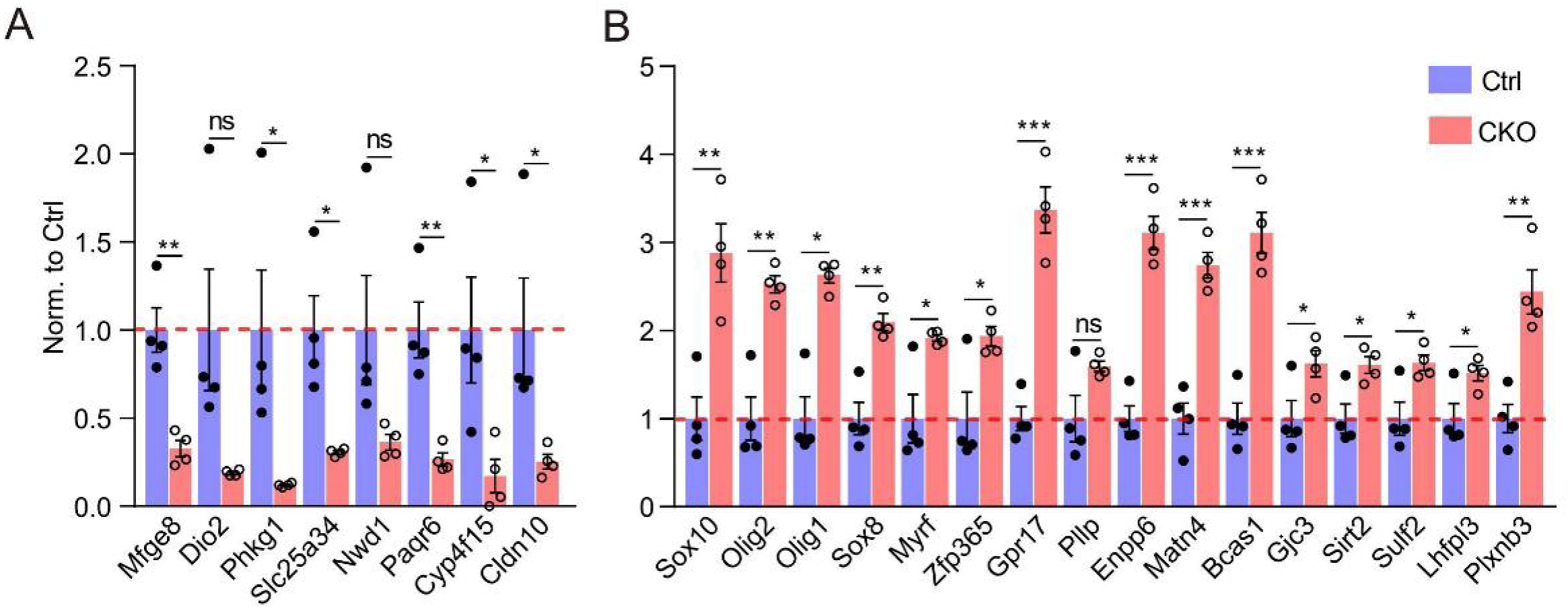
Alterations in gene expression pertaining to astrocytes and oligodendrocytes in P7 *_hGFAPAdam10-CKO_* _mice._ (**A**) Alterations in the relative abundance of astrocyte-related gene expression from the transcriptomes of P7 control (Ctrl) and conditional knockout (CKO) samples. (**B**) Alterations in the relative abundance of oligodendrocyte-related gene expression from the transcriptomes of P7 control and CKO samples. Control and CKO groups: n = 4 mice for each group. Two-tailed unpaired *t*-test: * *p* < 0.05, \*\**p* < 0.01, \*\*\**p* < 0.005; ns, not significant. Error bars, s.e.m.

**Supplementary figure 4.**
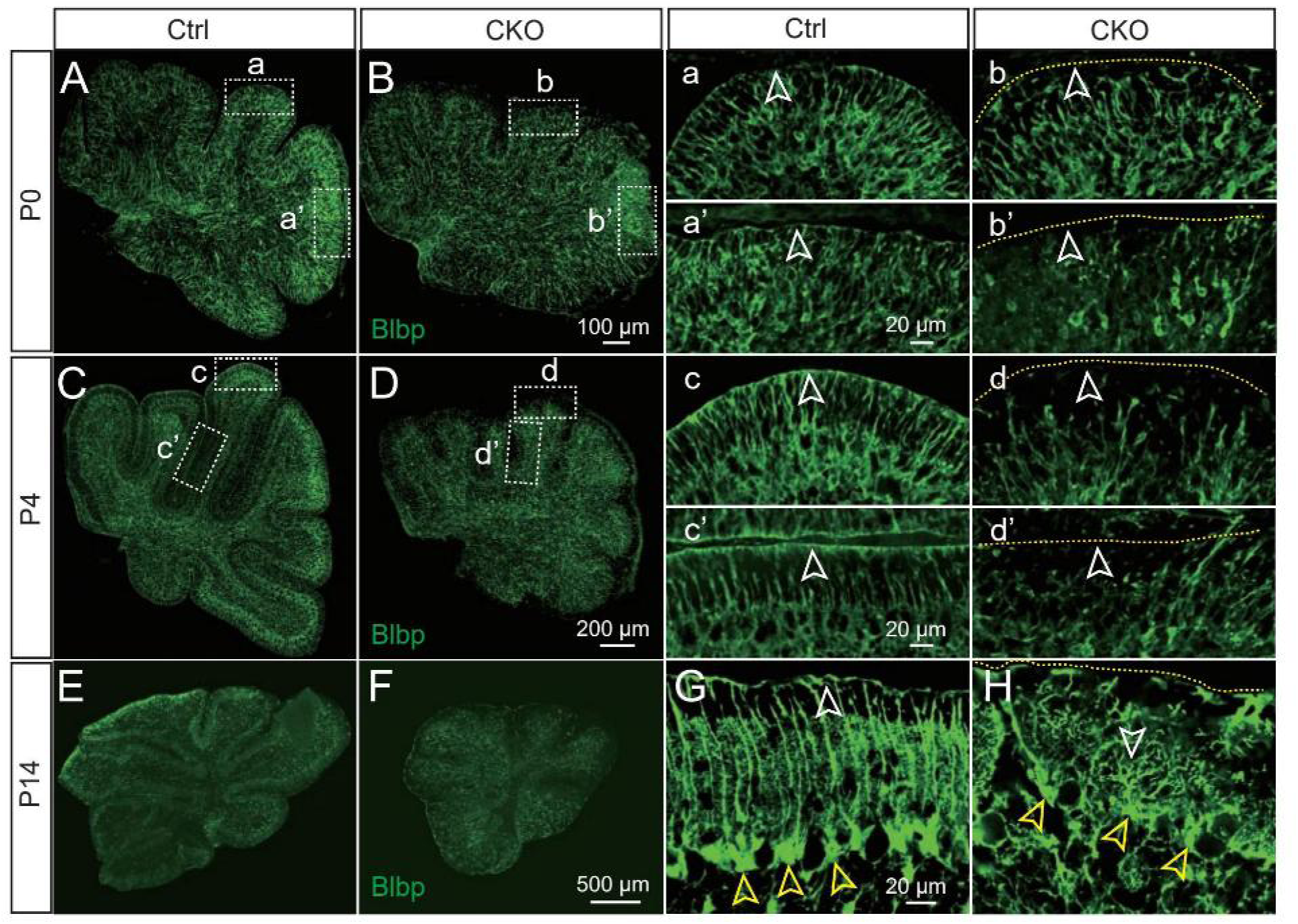
The morphology of astrocytes in the cerebellum of *hGFAP^Adam10-CKO^* mice. (**A–H**) Representative images of cerebellar sections stained with anti-Blbp, an antibody that recognizes astrocytes, in control (Ctrl; A, C, E, G) and *hGFAP-Cre;Adam10^fl/fl^* (B, D, F, H) mice. Three different developmental stages were analyzed: P0 (A, B), P4 (C, D), and P14 (E, F). (a–d) and (a’–d’) show the boxed areas in (A–D). The white and yellow open arrowheads, the pia of the cerebellum from control and CKO mice, respectively. The yellow dotted lines, the margin of the cerebellum in *hGFAP^Adam10-CKO^* mice.

**Supplementary figure 5.**
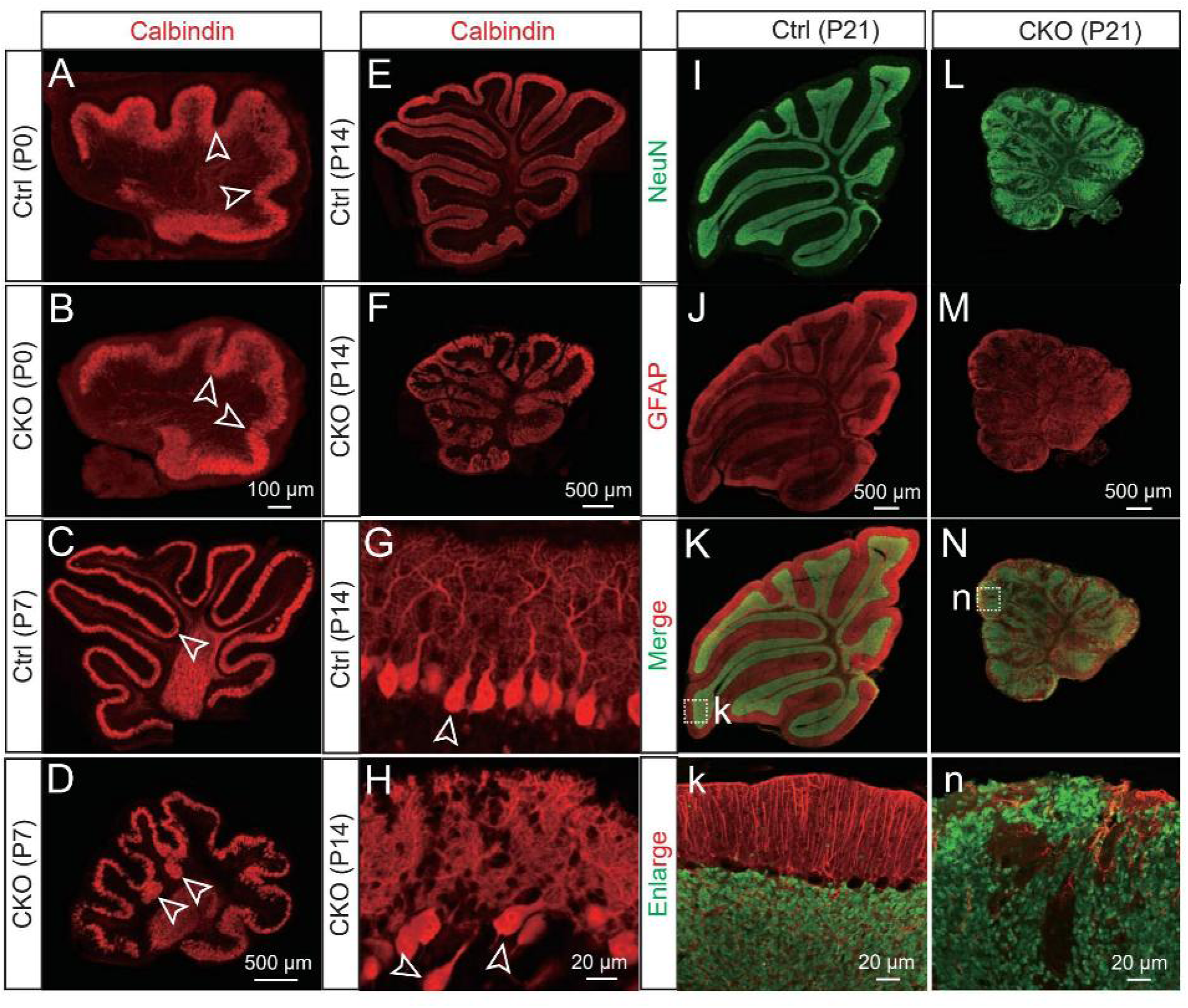
The morphology and distribution of neurons in the cerebellum of *hGFAP^Adam10-CKO^ mice*. (**A–H**) Representative images of cerebellar sections stained with anti-Calbindin, an antibody that recognizes Purkinje cells, from control (Ctrl; A, C, E, G) and *hGFAP^Adam10-CKO^* (CKO; B, D, F, H) mice at different developmental stages: P0 (A, B), P7 (C, D), and P14 (E–H). (**I–N**) Representative images of cerebellar sections stained with anti-NeuN and anti-GFAP to label neurons and astrocytes, respectively, from control (I–K) and *hGFAP^Adam10-CKO^* (L–N) mice at P21. (k, n) Higher magnification of the boxed areas in K and N, respectively.

**Supplementary figure 6.**
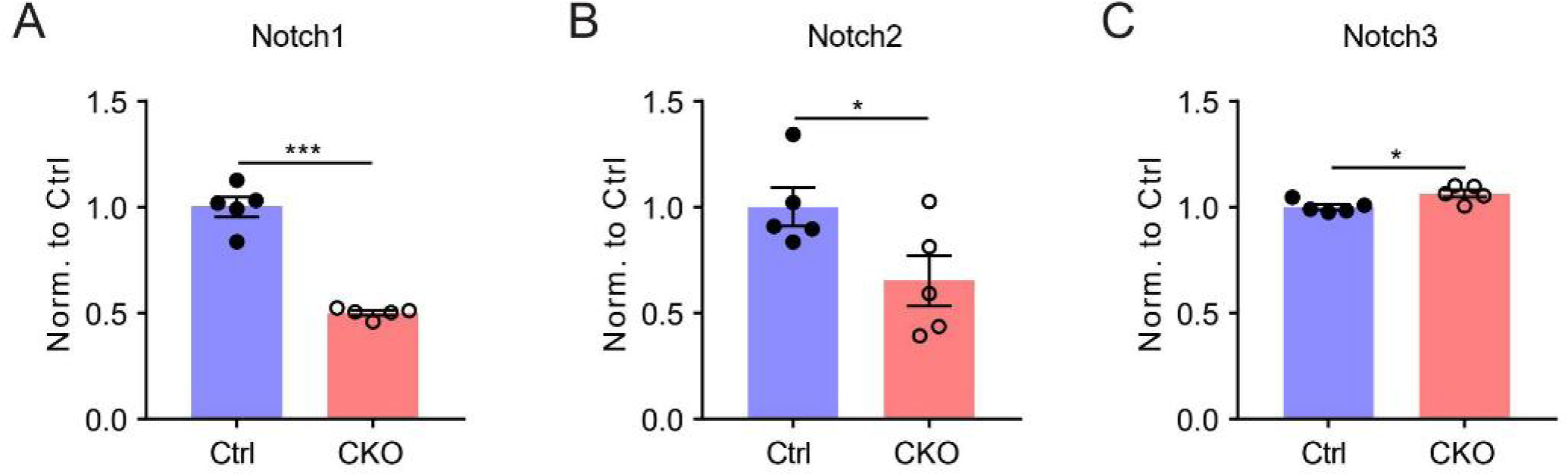
Alterations in gene expression of Notch1-3 in P0 hGFAPAdam10-CKO mice. (**A-C**) Alterations in the relative abundance of Notch1 (A), Notch2 (B), Notch3 (C) gene expression from the transcriptomes of P0 control (Ctrl) and conditional knockout (CKO) samples. Control and CKO groups: n = 5 mice for each group. Two-tailed unpaired *t*-test: * *p* < 0.05, \*\*\**p* < 0.005. Error bars, s.e.m.

## Author Contributions

W.G., S.D. W.W. conceived and supervised the project. Y.W., Y.L, and W.G. designed the experiments. Y.W., F.Z., X.C., X-F.G, and W.G. provided transgenic animals and performed animal experiments. Y.L. and T.Z. prepared cells for single-cell RNA seq. Y.W., J.-L. G., W.G. performed bulk RNA seq. D.A. analyzed data of single-cell RNA Seq. J. L. analyzed data of bulk RNA seq. Y L., Y. W., Z-B. G. completed staining experiments. W.G., S.D. W.W., L-J. W., W.S., S.Z., and Y.W. provided reagents. Y.L., Y.W., and W.G. wrote the manuscript. All authors discussed, reviewed, and edited the manuscript.

## Acknowledgment

We thank lab members at Ge lab from Chinese Institute for Brain Research, Beijing (CIBR), for their advice and feedback. xx for critical reading of the manuscript. We thank Drs. Qingchun Guo, Xinwei Gao, and Mingyue Jia from CIBR imaging core of CIBR for assistance in imaging and data analysis. We thank Drs. Wenlong Li, MS Ruirui Shen, and MS Shufang Huang at CIBR animal resource center for assistance in animal care. This work was supported by the Ministry of Science and Technology China Brain Initiative Grant to W.G (2022ZD0204703) and the Natural Science Foundation of China to W.G (32170964).

## Data availability

The datasets generated and analyzed in this study are available from the corresponding authors upon reasonable request.

## Methods

### Mice

All animal experiments were conducted following protocols approved by the Institutional Animal Care and Use Committee at Chinese Institute for Brain Research, Beijing (CIBR). *Ai14* (stock No. 007914), *hGFAP-Cre* (stock No. 004600), and *Adam10-flox* (stock No. 009357) were purchased from Jackson Laboratory. All mouse lines were genotyped as Jackson Laboratory recommended. We used the primers 5’-GAGAGGAAAGAAAGTGGCAGA-3’ and 5’- AGTGGGTGGGTTAATGAG CA-3’ for *Adam10-flox,* which gave a 370 bp band for the *Adam10* floxed allele and a 212 bp band for *the Adam10* wild-type allele.

### Behavioral tests-bar crossing tests

We performed the tests as previously reported(Clark et al., 1997). Briefly, a bar-crossing apparatus(Gerlai et al., 1993) was used to test balance and motor coordination. The device was a horizontal U-shaped platform supported on 30 cm tall legs. The two parallel bars of the U platform were 30 cm long and 18 mm wide, making it easy for mice to walk on. A narrow bar (30 cm long and 2 mm wide) connected the two wide bars, and was challenging for mice to walk on. Mice were placed on the wide bar at the beginning and moved freely for 10 min before following tests. In our tests, over the course of 2 min (120 s) of spontaneous activity on the wide bar, we evaluated the duration of motor activity (Locomotion Time), the duration of inactivity (Inactivity), the frequency of mice putting at least two feet away from the narrow bar and trying to cross to the wide bar (Cross attempts); the frequency of a mouse having one or more legs accidentally slipping off a wide bar (Slips); and the frequency of falling from the wide bar (Falls). In the 120-s measurement, we calculated the total durations of Locomotion, Inactivity, and Falls as 100%, as shown in Figures 1H and I. After completing the tests of spontaneous activity on the wide bar, mice were forced to cross the narrow bar. In this task, mice were placed on the middle of the narrow bar to force a crossing attempt. The longest time the mice stayed on the bar was 120 s, and the latency of successful crossing (Cross) or falling (Fall) was measured in seconds. A forced cross score was calculated as follows: if the cross was > 0, then forced cross score = 120-Cross; if the fall was < 0, then Forced Cross = Fall-120. Each animal was assigned a value between +120 (most agile) and −120 (least agile). Ethanol (70%) solution was used to clean the surface of the apparatus between tests with different mice. Mouse performances on the bar crossing apparatus were videotaped and manually scored as reported (Gerlai *et al*., 1993).

### Rotarod test

We used a rotarod test apparatus (Panlab, Model LE8205) to measure motor coordination after training. The initial speed of the setup was 4 rpm. The ramp-up time was 280 s. The maximum speed was 40 rpm. This experiment was conducted on mice 5–6 weeks of age. Mice were trained for three consecutive days before testing. The latency to fall was measured. Each mouse was scored three times/day for three consecutive days. The average score was recorded.

### Gait analysis

Mice were set to walk through a tunnel (length, 40 cm; width, 9 cm; height, 6 cm). The footprints were videotaped and analyzed. Step length was calculated as the average distance between one paw touching the ground and the opposite paw touching the ground. Step width was the average lateral distance between the opposite left and right steps. Stride length was the average distance between one paw touching the ground and the same paw touching the ground again. Stride time was the average time between one paw touching the ground and the same paw touching the ground again. The velocity was obtained by calculating the ratio of stride length to stride time to detect the motor ability of the mice. Furthermore, the change in mouse step width was determined by calculating the ratio of step width to stride length to exclude the influence of body size differences.

### Histology

Mice were perfused with phosphate-buffered saline (PBS) and 4% paraformaldehyde (PFA). Brain tissue was post-fixed in 4% PFA for 2h at 4℃, then stored in 30% sucrose solution in PBS before sectioning. Brain samples were sectioned by a cryostat (CM3050S, Leica). Sections with 5 μm thickness were used for hematoxylin and eosin (HE) staining. HE Staining Reagent Kit (Beyotime, Catalog C0105S) was used for staining. Briefly, samples were rinsed in de-ionized water for 2 min after air drying for 30 min. Next, sections were stained with hematoxylin for 10 min. After they were washed in tap water for 10 min, sections were washed once in de-ionized water. Next, sections were rinsed in 95% ethanol for 5s and then stained in eosin solution for 30 s. Next, they were washed in 70% ethanol twice, then in 95% ethanol twice for 5 min each, 100% ethanol twice for 5 min each, and dipped in xylene twice for 5 min each. Finally, sections were mounted with a resinous mounting medium.

### Bulk RNA-seq and Data Analysis

1. Sample preparation and RNA extraction P0 and P7 mouse pups were anesthetized with ice before decapitation. Cerebellums were collected for RNA extraction. Qiagen RNeasy Mini Kit (Qiagen, 74104) was used to extract the cerebellar RNA. SMARTer Stranded Total Sample prep kit-HI Mammalian (Takara, 634875) was used to generate libraries for RNA sequencing.
2. RNA sequencing (RNA-Seq) and differential expression analysis RNA-seq library construction for next-generation sequencing and paired-end deep sequencing were performed on an Illumina NextSeq500 SE75 platform (Illumina, San Diego, CA) according to the manufacturer’s protocol. Fastp(Chen et al., 2018) was recruited to low-quality reads and adaptor trimming with a default setting(Bolger et al., 2014). Cleaned reads were mapped to the Ensembl mouse reference genome GRCm38.p6 using STAR alignment software(Dobin et al., 2013). The mapped reads were counted to genes using featureCounts. Differential expression analysis was performed using DESeq2 with a cutoff of FDR < 0.05 and abs(log_2_FC) > 1. Principal component analysis (PCA) clustering plots, volcano plots, scatter plots, and heatmaps were generated using R packages (stats, ggplot2, and pheatmap) implemented in R Studio. In the heatmap plots, differentially expressed genes (DEGs) are grouped by hierarchical clustering analysis with parameters: clustering_distance_rows = “euclidean,” clustering_method = “ward.D2”.
3. GO and KEGG enrichment analysis Functional enrichment in GO terms (Cellular Component, Biological Process, Molecular Function) of differentially expressed genes (FDR<0.05 & |FC| >2) was performed using the clusterProfiler R package(Yu et al., 2012), setting a q value threshold of 0.05 for statistical significance.
4. GSEA analysis For gene set enrichment analysis (GSEA)(Mootha et al., 2003; Reimand et al., 2019; Subramanian et al., 2005), we manually organized genesets including immature astrocyte-enriched genes and mature astrocyte-enriched genes, OPC and pre-OL enriched genes, myelin OL-enriched genes(Cahoy et al., 2008) and a KEGG_2019_Mouse geneset based on database file from Enrichr online library(Chen et al., 2013; Kuleshov et al., 2016). Genes were pre-ranked through the metric algorithm (we applied sign of log fold change * -log10 (p-value[not adjusted p-val); the statistical result of DESeq2. Pre-ranked (.rnk) file and custom geneset were used as input for GSEA v4.2.2. The number of permutations was set at 1,000 and enrichment statistics were set to “weighted.” For general significance threshold, false discovery rate (FDR) q-val < 0.25 and |NES|>1.5 were considered significant enrichment.

### Single-cell RNA-seq and data analysis

Mice (P2) were anesthetized with ice before brain tissue was collected in ice-cold artificial cerebrospinal fluid (ACSF) containing 124 mM NaCl, 26 mM NaHCO_3_, 5 mM KCl, 2.6 mM CaCl_2_, 1.3 mM MgCl_2_, 1.25 mM NaH_2_PO_4_, 10 mM glucose, and 5 μM MK-801 at a pH of 7.4 when oxygenated (95% O_2_ and 5% CO_2_). Further dissection of the cerebellum was done under a stereoscope. Cerebellums were chopped with an aseptic surgical blade, and then the tissue was resuspended in ice-cold ACSF. Trypsin (5%, Fisher Scientific, Cat. No.: 15090046) and DNAse (2%, Sigma-Aldrich, Cat. No.: 10104159007) were added into the resuspension and placed into a pre-heated 37℃ water bath for 10 min. After 10 min, a sterilized glass pipette was used to blow gently 15–20 times to help digestion. FBS was added (10%, Fisher Scientific, Cat No.: 10-099-141) to stop digestion, and 10% OptiPrep (Sigma-Aldrich, Cat No.: D1556-250ML) was added for gradient centrifugation at 4℃ (900 g, 10 min). We then removed the supernatant with a pipette and washed it with ACSF before centrifuging again (900 g, 3 min, 4℃). Next, we removed the supernatant and added 1 ml ACSF with DAPI (Vector Laboratories, Cat No.: H-1200-10ml, 5 μ g/ml). Collected single-cell suspensions were used for library construction.

The count matrix of a single cell was preprocessed by Cellranger (V3.1.0) with the reference genome (Version: GRCm38). The downstream analysis was performed with Scanpy and its extending packages. Low-quality single cells were filtered out, with the number of expressed genes in a cell over 200 and the number of cells expressing the gene over 3. Doublet detection was performed with the extending package SOLO from scvi-tool. The percentage of counts in mitochondrial genes was below 0.1, and the number of genes expressed in the count matrix was below 6,000. Merging single-cell datasets was performed with Harmony. The embedding of a single cell was performed with UMAP [<u>10</u>]. The clustering of cell subpopulations was performed with Louvain. The marker genes were found by Wilcoxon rank-sum test with corrected ties. The RNA velocity was generated from Velocyto. Finally, the cell fate analysis was performed with Cellranger.

### Immunostaining

We sectioned fixed cerebellums with a thickness of 30 μm. Brain sections were blocked in buffer with 0.2% Triton-X-100, 3% BSA (Sigma, A4503), and 5% Normal Goat Serum (Jackson Immuno Research) for 2 h at room temperature. After blocking, sections were incubated overnight with primary antibodies for NeuN (1:100, Chemicon, mab377), Calbindin (1:3,000, Sigma, C2724), Blbp (1:1,000, Sigma, ABN14), GFAP (1:300, Sigma, G9269), Olig2 (1:100, Abcam, ab109186), or MBP (1:1,000, BioLegend, SMI-99P-100). After sections were washed in PBS 3 times for 10 min each, secondary antibodies Alexa Fluor 488 goat anti-mouse (1:500, Thermo, A11001) or Alexa Fluor 488 goat anti-Rabbit (1:1,000, Thermo, A11008) were incubated with Hoechst 333342 for 2h at RT.

